# Play it by Ear: A perceptual algorithm for autonomous melodious piano playing with a bio-inspired robotic hand

**DOI:** 10.1101/2024.06.04.597044

**Authors:** Hesam Azadjou, Ali Marjaninejad, Francisco J Valero-Cuevas

## Abstract

Perception shapes the learning and performance of motor behavior in animals. In contrast to this inherent biological and psychological connection between perception and action, traditional artificial intelligence methods for robotics emphasize reward-driven extensive trial-and-error or error-driven control techniques.

Our study goes back to the perceptual roots of biological learning and behavior, and demonstrates a novel end-to-end perceptual experience-driven approach for autonomous piano playing. Our ‘Play it by Ear’ perceptual learning algorithm, coupled to a bio-inspired 4-finger robotic hand, can replicate melodies on a keyboard after hearing them once—without explicit or prior knowledge of notes, the hand, or the keyboard. Our key innovation is an end-to-end pipeline that, after a brief period of ‘motor babbling’ by the hand, converts the sound of a melody into native musical percepts (note sequences and intensities) that it replays as sequences of key presses.

In this way, any new melody consisting of notes experienced during babbling can be reproduced by the robotic musician hand on the basis of its percepts. This playback includes capturing the qualitative and quantitative musical dynamics and tempo with a nuance comparable with that of four human pianists performing the same melody. These compelling results emphasize the perceptual underpinnings of artistic performance as an alternative to traditional control-theoretical emphasis on state estimation and error correction. Our approach opens avenues for the development of simple machines that can still execute artistic and physical tasks that approach the nuance inherent in human behavior.

## Main

In human development and artistic expression, motor behavior is intricately intertwined with perception(1; 2; 3; 4; 5). This fundamental notion is well-established in psychological and psychophysical studies. In contrast, traditional robotics relies heavily on control theory, extensive trial-and-error, and precise models of the physical system, task, and/or environment. Alternatively, expert demonstration of the task or expert knowledge are used to fine-tune these systems which often require extensive training in simulation or via interactions with the environment(6; 7; 8; 9; 10; 11; 12; 13; 14).

We demonstrate it is possible and efficient to emulate the human ability to replay a melody after hearing it by developing a bio-inspired perceptual end-to-end pipeline. As proof-of-principle of robotic artistic expression, we developed a novel a (*musician hand*) that can reproduce a melody on a keyboard driven solely by the perception of the music. Rather than focusing on the mere detection and control of finger movements, or mapping Instrument Digital Interface (MIDI) notes to finger motions, our system replicated the perceptual essence of the music (note sequences and intensities), akin to the intuitive and experience-driven nature of human *‘playing by ear’*(15; 16; 17; 18).

Perception plays a critical role in shaping the coordination, stability, and organization of motor actions (19; 20; 21; 22; 23; 24; 25; 26; 27; 28; 29; 30). Perception fundamentally contextualizes and attributes meaning to sensations to produce meaningful actions. It is a process built upon experience and interactions with the physical world that creates a *Sensorimotor Gestalt* of compatible sensory sets and motor actions (i.e., the perception-action link) (31; 32; 33; 34) These ideas have been proposed as potential foundations of Identity, Agency, and Self for robots (35). From a computational perspective, perception is also implicit in the nature of reinforcement learning: A sensory signal, to be useful in learning, is often imbued with the meaning of ‘fitness,’ ‘value,’ ‘reward,’ or ‘cost’—which are used by control policies to learn to take appropriate actions that are ‘reinforced,’ learned and exploited. Recent research highlights the promise of such experience-driven methodologies for reinforcement and imitation learning to dynamically learn and adapt via interactions with the physical world. (36; 37; 38; 39; 40; 41; 42) but, importantly, they are not in the domain of artistic performance by robots.

We translate end-to-end autonomous learning architectures to the perceptual domain of experience-driven artistic performance. The value of sequences of physical actions lies in their melodic nuance. As a first example, we used piano playing as it involves note sequences and intensities that are woven together via delicate control of the timing and dynamics of finger actions to elicit a melodious experience in the listener.

Importantly, this work is a departure from XIX century pianolas, or modern physical or simulated robotic hands that can play the piano (43; 44; 45; 46; 47). Instead of driving the musical performance by punch cards, magnetic tape or Musical MIDI code that spells out the melody, our approach is to drive melodious performance from the perception of melodies themselves—as human pianists can. Moreover, our approach (Fig. 1 and 2) leverages the biological advantages of brain-body co-evolution and co-adaptation (48; 49). The physical properties of our robotic hand were specifically adopted to contribute to the nuance of the music played by using a tendon-driven design actuated with DC motors emulating muscle contraction (Fig. 3) (50), and foam-covered fingertips to approximate the compression of the finger pads of human pianists (51) (Fig. 3). Thus we pioneer a distinctive approach employing end-to-end bio-inspired one-shot learning, which we call *‘Play it by Ear’* (Fig. 1).

**Figure 1.**
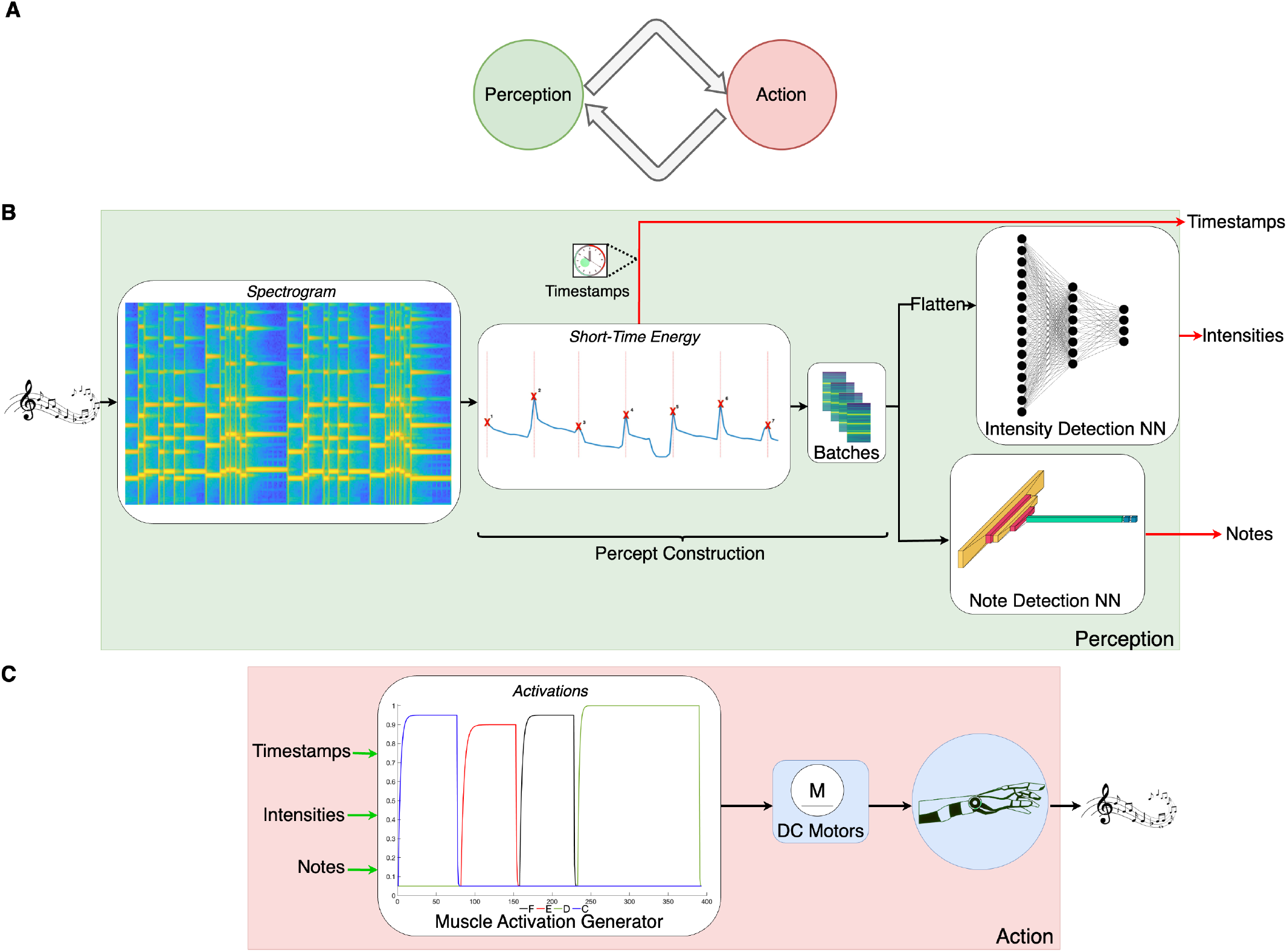
Overview of the ‘Play it by Ear’ algorithm. **A**. The perception-action loop is a fundamental principle of biological behavior, especially in artistic expression. **B**. Our algorithm takes a melody (sensed as a spectrogram) and extracts batches of ‘percepts’ (sequences of notes and their intensity) on the basis of prior ‘motor babbling’ experience (see Figure 2). **C**. The melody is replicated by playing the notes on an actual keyboard by the four fingers of a bio-inspired robotic hand.

**Figure 2.**
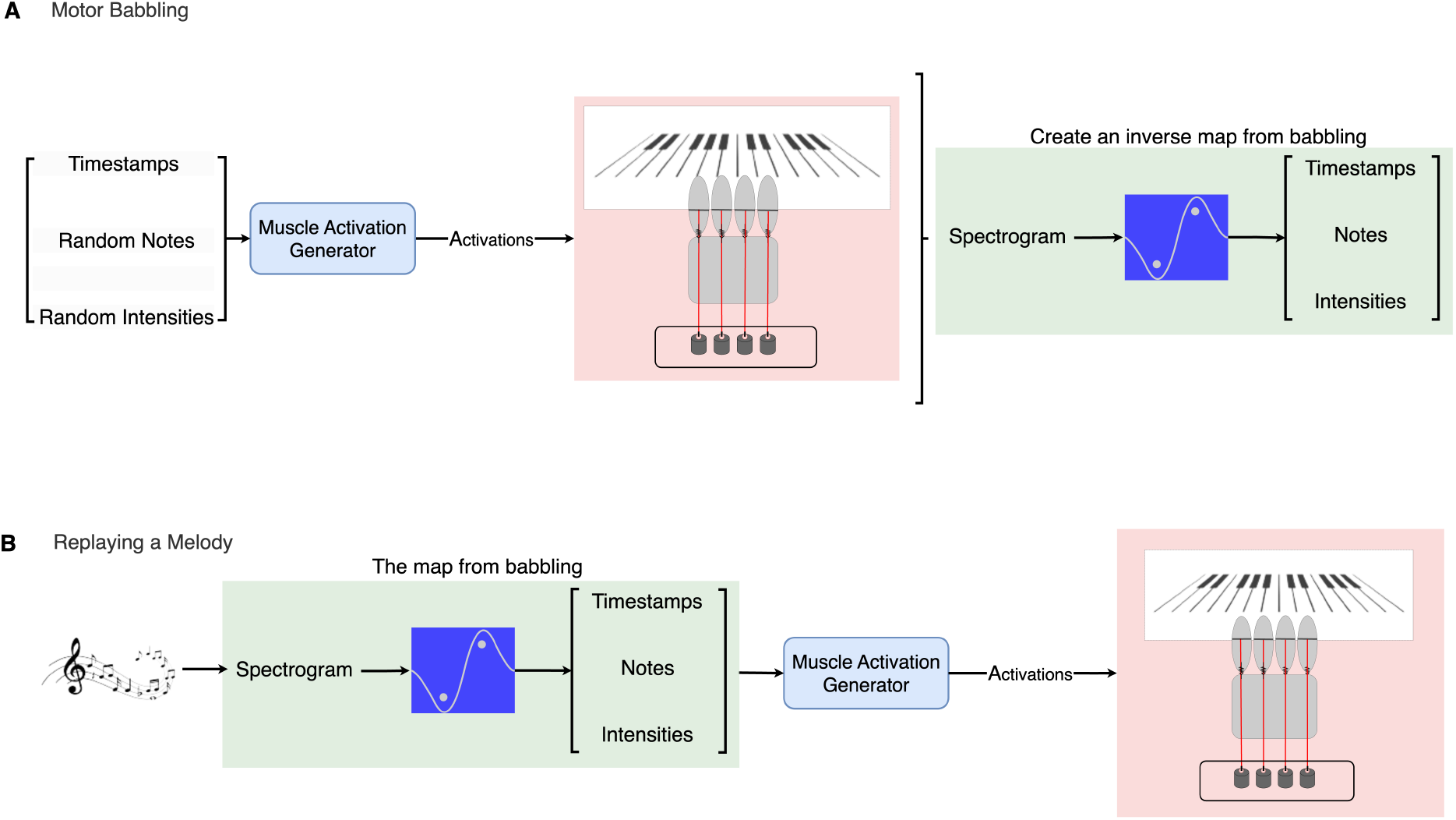
Motor babbling and how it is used to play a melody ‘by ear.’ **A**. The algorithm starts from a naïve state and, like a novice human, randomly explores the mapping from finger actions to percepts (i.e., note sequences and intensities). We implemented such ‘motor babbling’ by recording 2 mins. of random sequences of individual finger actions, each lasting 500 ms. We used these input-output pairings to train artificial neural networks (ANNs) to detect percepts, to then produce finger actions that replicate the melody. **B**. This allows the algorithm to command the robotic hand to play any arbitrary melody that includes the notes experienced on the keyboard.

**Figure 3.**
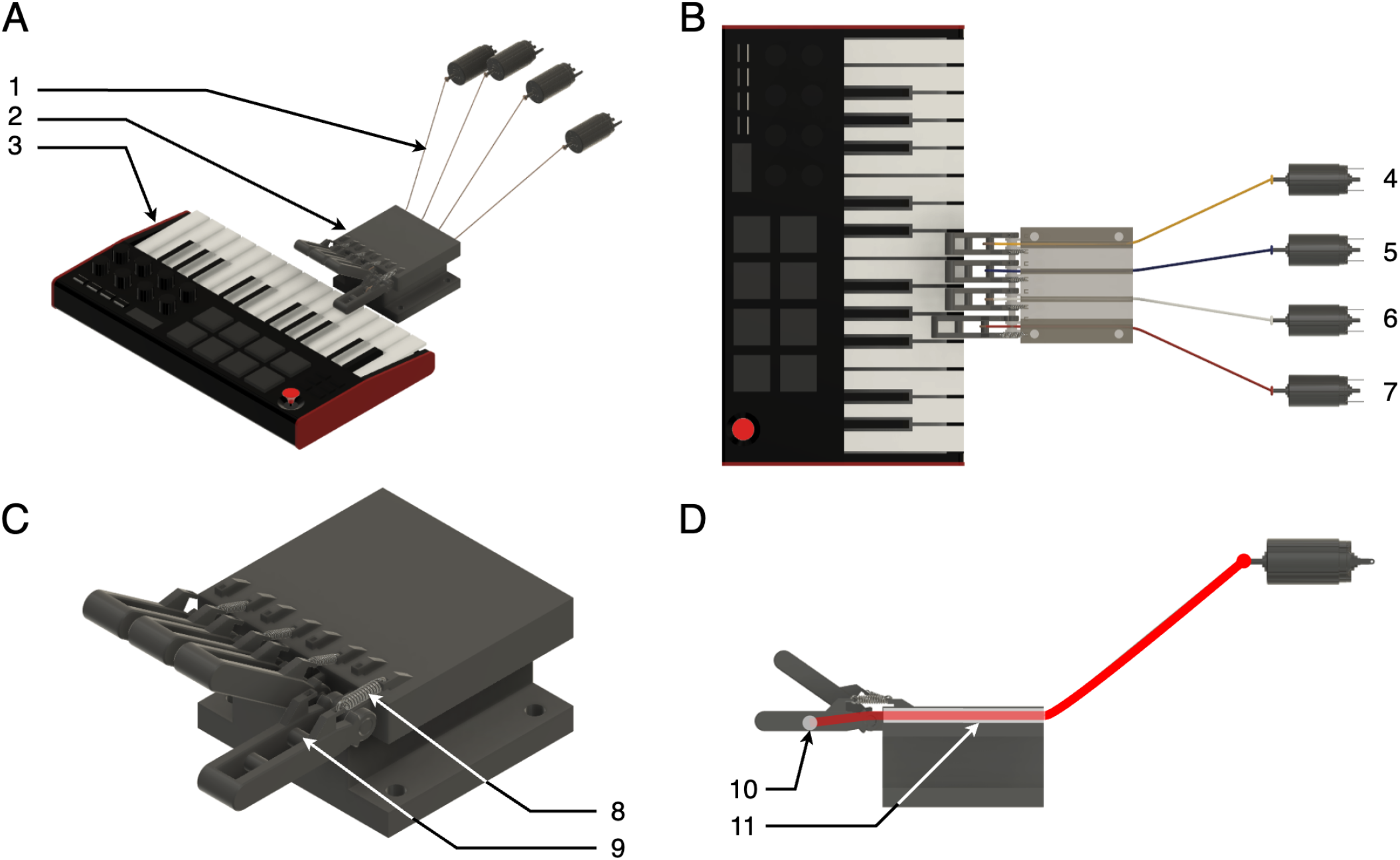
A bio-inspired robotic hand uses four fingers to perform motor babbling to then play melodies by ear. **A**. Overview: 1. Flexor tendons pulled by DC motors. 2. Hand’s structure. 3. Keyboard. **B**. Top view of the system: 4–7. the four DC motors (Motor0–3) used to pull the tendons. **C**. Hand: 8. Spring to passively extend finger. 9. Flexor tendon attachment. **D** Side view of the hand: 10. Flexor tendon attachment 11. Flexor tendon routing.

Our *‘Play it by Ear’* approach emulates human learning by starting with ‘motor babbling’ to train the perception-action map. We do this by first commanding pseudo-random sets of motor activations to each finger while recording their musical output as sound (Figure 2A). This allows us to create the inverse map (perception-action strategy) that finds the motor activations needed to replay any melody of interest by detecting the musical percepts of notes and their intensity (Figure 2B).

Our findings demonstrate that we can leverage the foundations of the perception-action loop to create end-to-end robotic systems that can produce melodious artistic expression on the basis of limited experience—and without explicit knowledge on the task, system model, and state variables (i.e., agnostic to the specific nature of the hand, musical instrument, or style of the music).

## Results

We show that using perceptual learning (PL), the musician hand learns to replay any arbitrary 4-note melody without closed-loop error correction or an explicit model of the dynamics of the tendon-driven fingers or the physical environment (e.g., keyboard inertia or contact dynamics) in a one-shot learning manner. We also show how the musician hand can outperform novice participants and replays a melody on comparable level with trained human pianists. The PL of the musician hand is inspired by a fundamental principle embedded within human biology—the perception-action loop. To replicate this loop in our Play it by Ear algorithm, it receives the perceptual inputs (batches of spectrograms) and processes them into motor-action commands to run the motors and replay the target melody.

### Perceptual Learning

To train the musician hand in replaying a melody from scratch, we employed motor babbling to learn how to map sensory information (sound) to motor actions (notes played). A run from a na ive state started with 2 mins. of babbling, where a sequence of pseudo-random motor activations (using activation profiles inspired by Hill-type muscle models, Fig. 4) were distributed across the four motors controlling the tendons. Each motor action lasted 500 ms. and pressed keys at random with varying forces to produce notes of different intensities, Fig. 2A. We converted the resulting 2-min. audio signal into a spectrogram using Short-Time Fourier Transform (STFT). The spectrogram is the perceptual cornerstone to train two ANNs to associate note sequences and intensities from sound (i.e., percepts) with motor actions. These ANNs then are employed to predict notes and intensities when hearing a new melody. One of the ANNs is a Convolutional Neural Network (CNN) trained to detect the notes played at each percept—using a 200 ms interval after note onset that assumes a ^4^ *andante* tempo. To capture the nuances of the melody and replicate its musical dynamics, the second ANN (a Multi-Layer Perceptron, MLP) was trained to map the percepts into four levels of intensity (i.e., varying degrees of loudness). The validation of the CNN and MLP trained on babbling data is detailed in Table 1. The last musical element, note onset, was automated through the identification of peaks in the short-time energy function of the audio signal derived from the target melody’s spectrogram (see Fig. 5).

**Table 1.** Validation Accuracy of the Networks on Babbling Data.

| Network | Cross-Validated Accuracy (%) |
| --- | --- |
| Note Detector Network | 98.4 |
| Intensity Estimator Network | 86.2 |

**Figure 4.**
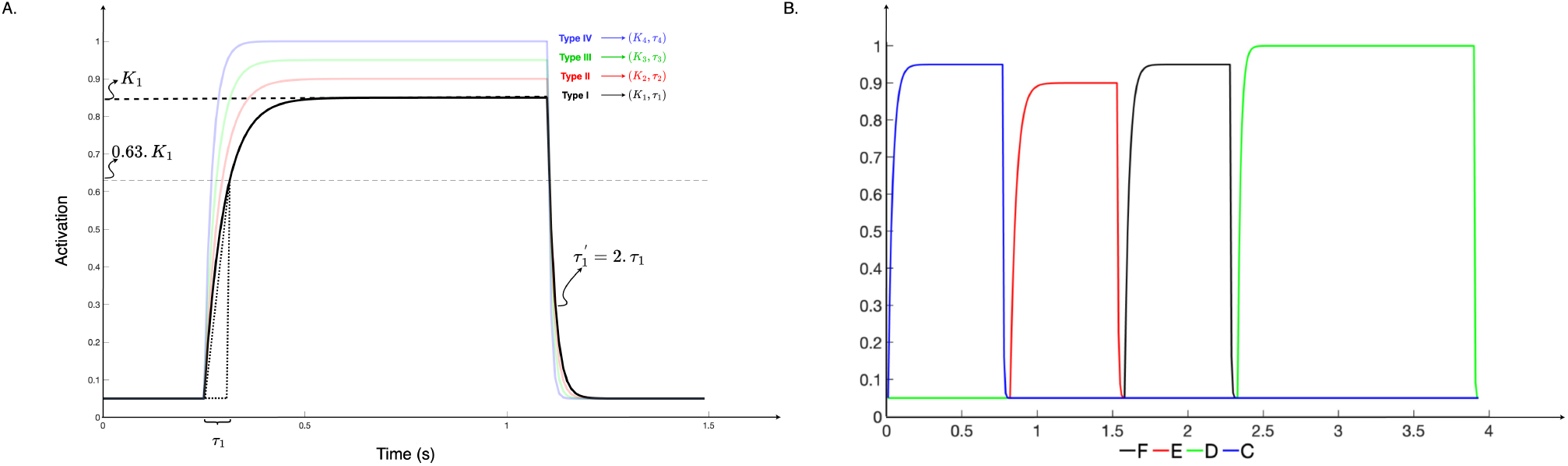
A. The four motor activation profiles used by the musician hand. We pre-programmed four activation profiles to replay the melodies. The exponential rise and fall are inspired by Hill-type muscle model (50; 52), and reached four different intensity plateaus, *K*_1−4_, and time constants, *τ*_1−4_. This produced four different key press mechanics leading to different note onset, loudness and decay.**B. Motor Activation Sequences:** In this figure we see an example of the activation sequence sent to the motors to pull the tendons and press the keys.

**Figure 5.**
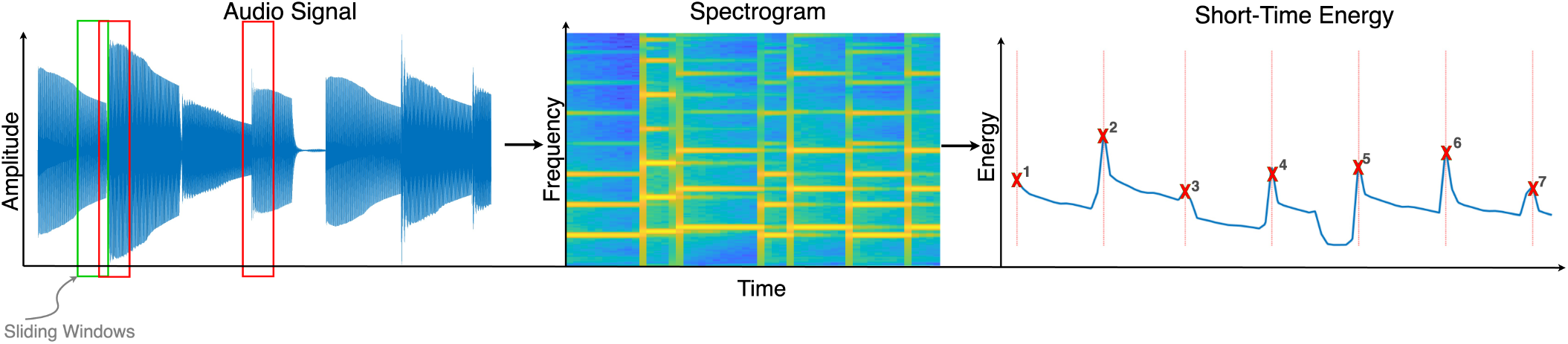
Signal processing transforms the percept of a melody into signals that can train the ANNs. As shown in Fig. 1B, we extract the time-frequency representation (spectrogram) of the audio signals using sliding windows that compute the Fourier transform of the signal in short overlapping time intervals. As STFT computes the signal’s power in different frequencies for short time intervals, we integrate the spectrogram on the frequency axis to compute the short-time energy (STE) of the audio signals that represent the notes onsets as their peaks (as you see the red windows that cover the second and the third note’s onsets are the second and third peaks in STE)

### Quantitative Results

The *quantitative* effectiveness of both the PL algorithm and human participants in replicating Melody 1 (M1) is demonstrated by the accuracy of note detection, intensity estimation, and timing, as illustrated in Fig.6A. All five novice and four human pianists read, understood and signed an informed consent form approved by the Institutional Review Board for the protection of human subjects at the University of Southern California. Human participants were allowed to babble for 5 mins. on the relevant keys (without shifting their fingers) and had three minutes to practice M1 before being given one minute to play the 29-second melody (i.e., ‘final performance’). To eliminate the need to memorize M1, they could navigate M1 forward and backward during the practice and performance periods. The five novice human players could only repeat the first two or three notes correctly, and none succeeded in playing the melody beyond the third note in the 1 min. provided—thus they had null results for Intensity Estimation and Time Difference (i.e., note missing red bars for these two metrics). On the other hand, all human pianists successfully reproduced the melody (one played flawlessly, while the others made one—four mistakes when performing the 37 notes) with very good Intensity Estimation (higher is better) and Time Difference (lower is better). By comparison, the human pianists slightly outperformed the musician hand, which had comparable quantitative scores for M1 (Fig. 6A). We also presented two new melodies (M2 & M3 in Fig. 7) to the musician hand without additional babbling or training, which it performed similarly well (additional blue bars in Fig. 6A).

**Figure 6.**
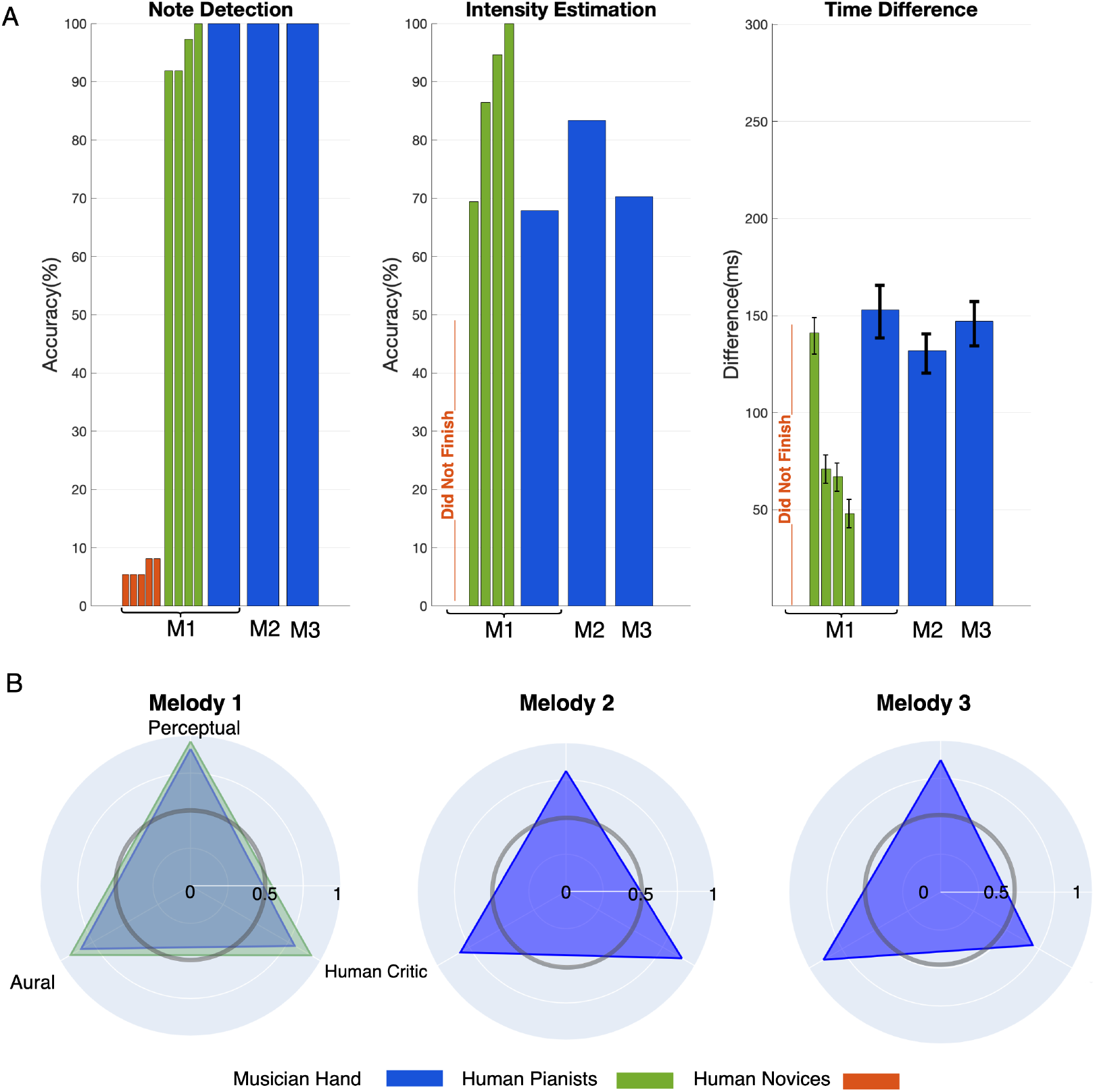
A. Quantitative Performance Results: While 5 human novices stumbled (red bars), and 4 human pianists performed well (green bars) for Melody 1 (M1), our musician had perfect Note Detection for three melodies (blue bars, M1–M3). Intensity Estimation (a score for replicating musical dynamics) was score-less for the human novices (i.e., they failed to replicate the melody). Human pianists scored between 78% and 100%, while the musician hand scored between 68% and 84%. Time Difference (smaller is better) quantified the replication of note onset and reflects rhythm across the melody. Human pianists scored best, at below 120 ms. The musician hand scored between 130 and 260 ms depending on melody. Error bars are standard deviations across all notes. **B. Qualitative Performance Results:** For each melody, we evaluated how melodious the replication was along three qualitative dimensions: **Perceptual** As per the structural similarity index metric between the spectrogram images (53). **Human Critic** As per the blinded rating by a professional pianist. **Aural** As per similarity of Mel-Frequency Cepstral Coefficients (approximating the human auditory system’s response (54)). For Melody 1, the human novices were patently worse, but the musician hand performed within 1 standard deviation of the human pianists along all three dimensions. The musician hand performed similarly for the other two melodies, which the human pianists did not play.

**Figure 7.**
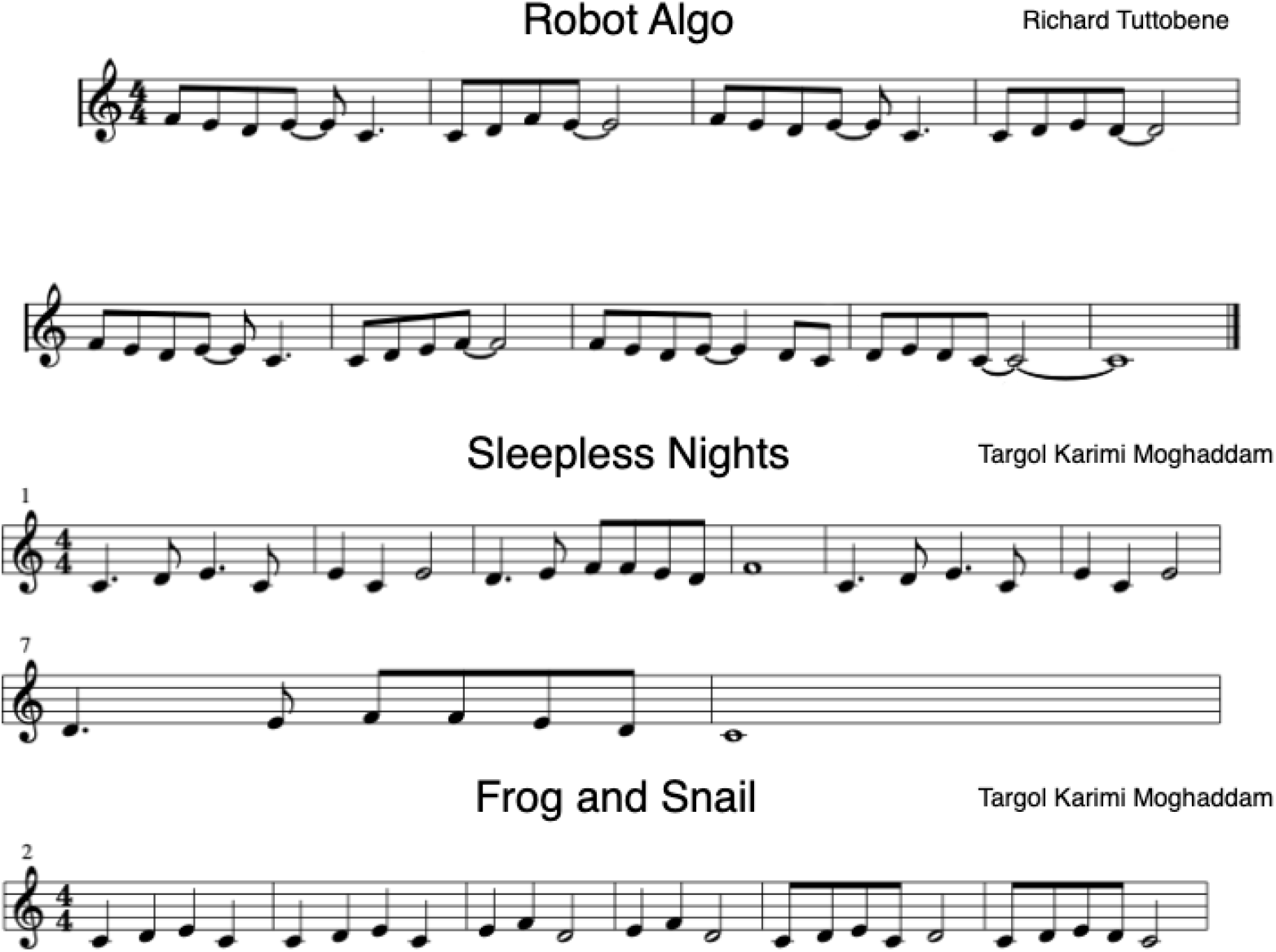
Scores for all three melodies: *M1: Robot Algo, M2: Sleepless Nights*, and *M3: Frog and Snail*.

### Qualitative Results

We also evaluated the *qualitative* performance of the recordings of M1 from the human participants and the PL algorithm. We did this along the three dimensions of Perceptual Similarity (as per the Structural Similarity Index (SSIM) of their spectrograms), a human critic’s rating, and Aural Similarity (cosine similarity of the Mel-Frequency Cepstral Coefficients, MFCCs), Fig.6B. This Figure shows that the PL algorithm was practically equal to the qualitative scores from the four human pianists. The human novices scored much lower than either. Note that the PL algorithm was also able to perform well in M2 and M3.

The SSIM of the spectrogram serves as a perception-based measurement, assessing the perceived alteration in structural data. It integrates key perceptual factors like luminance masking and contrast masking terms to measure the similarity between melodies’ time-frequency representations derived from their spectrograms (53). We also asked the human expert to rate the similarity of the melodies from 1 to 10 and normalized that to qualitatively assess the performance from a human critic’s perspective. Finally, we extracted the MFCCs, which represent the short-term power spectrum of a sound, to compare the cepstral representation of the audio clip (a nonlinear “spectrum-of-a-spectrum”). MFCC captures essential information, particularly low-frequency characteristics, more accurately than linearly spaced frequency bands, closely approximating the human auditory system’s response (54). The spider plot in Fig.6B shows the similarities.

## Discussion

Our study goes back to the perceptual roots of biological behavior to demonstrate a novel end-to-end perceptual experience-driven approach for autonomous melodious performance. As proof-of-principle, this ‘Play it by Ear’ perceptual learning algorithm using a bio-inspired 4-finger robotic hand to replicate melodies on a keyboard after hearing them once, without any prior information of the task, any explicit knowledge of the notes, hand, or keyboard states. Demonstrating a perceptual learning algorithm on hardware and using a bio-inspired hand (tendon-driven and activated by the Hill-type muscle model) unveils novel possibilities for bridging the gap between human-like perception and robotic autonomy.

Our PL algorithm can, without any prior knowledge and only by performing a brief period of ‘motor babbling,’ interpret (i.e., classify) the sound of a melody into musical percepts (note sequences and intensities). This enabled a robotic hand to artistically replay arbitrary melodies of any duration that use the notes experienced during babbling—in a single attempt and after hearing them only once. Importantly, the algorithm’s ability to ‘Play by Ear’ was, respectively, better than and comparable to novice and trained human pianist. This performance comparison involved quantitative and qualitative metrics (Fig. 6). It is likely that our implementation of the algorithm and hardware hand could not match the perceptual and/or motor abilities of trained human pianists, who train for years to excel in perceiving and playing music on the piano. Nevertheless, our findings showcasing human-like performance do demonstrate the feasibility and promise of developing and integrating perceptual algorithms with robotic hardware.

It is important to note that the robotic hand successfully replayed the target melody after limited prior experience (2 minutes of motor babbling) and hearing it only once. This is an example of one-shot learning (by not requiring or allowing refinements as defined in the field (55))—-which is moreover accomplished without explicit or prior knowledge of notes, the hand, the keyboard, or the target melody. In contrast, state-of-the-art robotic systems typically rely on extensive pre-programmed knowledge and feedback mechanisms during repeated attempts to refine a pre-defined known task (6; 8; 9; 10; 11; 12; 13; 56). Our approach, therefore, fills a significant gap in the field by demonstrating that a robotic system can achieve one-shot nuanced musical performance using minimal prior information and training.

The limitations of this first proof-of-principle demonstration point to several avenues of technical improvement and conceptual interpretations and speculations. First and foremost, the performance of the PL algorithm is—as in biological motor learning—a product of its own limited motor babbling experience with a finite set of intensities (Fig. 4. The experience of the PL algorithm is, by definition, a smaller set of musical perception and motor actions than the trained human pianists. This limited repertoire of experience, as in biology, naturally limits how the melodies were *perceived* and *interpreted* (i.e., heard and reproduced, respectively). This perceptual limitation of the PL algorithm is analogous to biological discretization of human hearing—whereby the perceptual field is not continuous, but rather a collection of attractors around the phonemes common in a mother tongue (57; 58). Thus, some spoken accents in a second language cannot be improved upon as the speaker simply cannot distinguish phonemes in the second language that fall into nearby attractors in the mother tongue. Thus, one source of inaccuracy could be the inability of the PL algorithm to detect nuance in the target melody that was not present in its limited motor babbling.

Regardless of these limitations, our work represents a departure from traditional artificial intelligence methods for robotics, which emphasize reward-driven extensive trial-and-error or error-driven refinement techniques. By going back to the perceptual roots of biological learning and behavior, we demonstrate a novel end-to-end perceptual experience-driven approach for autonomous piano playing.

This demonstration of the PL algorithm suggests it is possible to replicate human artistic behavior that inspire new forms of human-robot interaction and collaboration that approximate perception-driven (and at times emotionally-driven (59)) strategies that humans depend on (19; 20; 21; 22; 23; 24; 25; 26; 27; 28; 29; 30). Integrating advancements in artificial intelligence and biological principles could facilitate even more seamless human-robot collaboration, driving innovation in various domains (60). Additionally, incorporating real-time feedback mechanisms and adaptive learning could further improve the precision and expressiveness of robotic performances.

Future work could explore the adaptation of our algorithm to different types of musical instruments and performance styles, enhancing its versatility. A key element of musical performance in humans is our ability to listen to a piece of music played on one instrument and replicate it on a different one. This is a form of biological transfer learning (based on compositionality) that ANNs currently lack (60). This use of compositionality—the ability to decompose complex tasks into more elementary components that can be reused for related tasks (60)—would be particularly useful as an extension of our PL algorithm for musical performance. This fundamental limitation of ANNs comes from the fact that the training set has an inordinate influence on the setting of weights that somehow reduces the neural network’s capacity to generalize across varying instrumental styles (61). Future work should develop more advanced transfer learning techniques, potentially using perception help bridge the gap between learning algorithms and human artistic performance.

Lastly, expanding the PL framework to other sensory, visual and proprioceptive modalities could open new avenues for research and application in bio-inspired robotics and machine intelligence to, say, visual arts or acrobatic maneuvers.

## Methods

### Dynamic Control of Tendon-Driven Fingers

Our system controls four tendon-driven fingers to play piano keys using four DC motors. Each motor pulls a tendon, adjusting the corresponding finger’s position and velocity. The control strategy dynamically activates the motors based on the output of an artificial neural network (ANN) that classifies musical notes and their intensities at different time points. Each finger *F*_*i*_ is modeled as a second-order system:

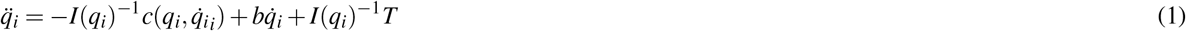

where 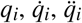 are the joint angle and its first and second dreivative respectively in finger i, *I* is the inertia,*c* is the Coriolis and centripetal force,*b* is the joint friction coefficient, *k* is the stiffness, and *T*_*i*_(*t*) is the torque in finger i generated by *M*_*i*_. The musculotendon forces (represented here as cables pulled by the motors) are subsequently correlated with the vector of applied joint torques:

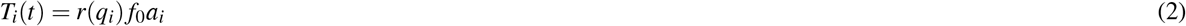

where *r* is the moment arm of each finger, *F*_0_ is the maximal force exerted by each motor and *a* is the normalized actuation of the motor. In order to predict the actuation values of each motor to play the melody, we use a mapping that connect a melody’s spectrogram to actuation time series:

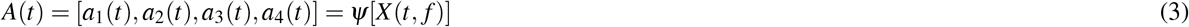

where *ψ* is a mapping consisting of an energy function, a CNN, an MLP and a motor activation generator:

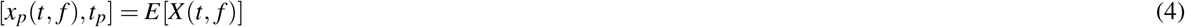

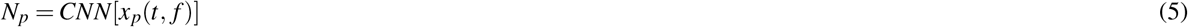

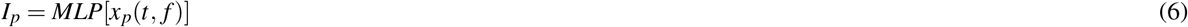

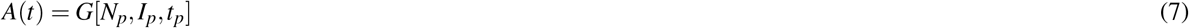

Where E is the energy function that extracts the timestamps for the note onsets (*t*_*p*_) and batches of spectrograms that start from the timestamps (*x*_*p*_). CNN and ANN then use these batches to extract notes and their intensities (*N*_*p*_,*I*_*p*_) and the motor function generater uses a Hill-type muscle activation profile (see Fig. 4) to generate a motor activation sequence that leads to the musician hand play the melody. We used four different intensity plateaus, *K*_1−4_, and time constants, *τ*_1−4_. This produced four different key press mechanics leading to different note onset, loudness and decay. The mapping adjusts the voltage applied to each motor, ensuring the fingers achieve the desired movements and note intensities in real time, as dictated by the percepts from the ANN.

### Time-Frequency Representation

We employed the Short-Time Fourier Transform (STFT) to extract spectrograms from the audio signals of the melodies under investigation. The STFT is a widely utilized signal processing technique for analyzing the frequency content of non-stationary signals over short time intervals. First, each audio signal *x*(*t*) was divided into short overlapping segments. Let *x*_*n*_(*t*) represent the *n*-th segment of the audio signal. The Fourier Transform was then applied to each segment to transform it from the time domain to the frequency domain. This process is mathematically defined as:

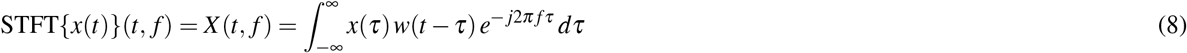

where *w*(*t*) is a window function that is non-zero for a short duration and *X* (*t, f*) represents the STFT of the signal *x*(*t*), giving the frequency content of the signal at time *t*. Repeating this process for all segments and stacking the resulting frequency spectra over time, we extracted a time-frequency representation of the melody’s audio signal, commonly known as a spectrogram. The spectrogram *S*(*t, f*) is given by:

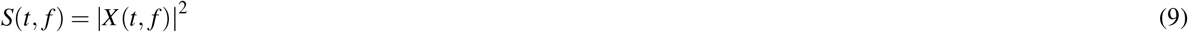

This spectrogram provided valuable insight into the distribution of frequencies present in the melody at different points in time. We used the spectrogram to extract the notes and their intensities to automate the musician hand (Fig. 5).

### Energy Function

We integrated the spectrograms along the frequency axis to derive the energy function of each melody. Let *S*(*t, f*) represent the spectrogram of the audio signal, where *t* denotes time and *f* denotes frequency. The energy function *E*(*t*) at time *t* is obtained by integrating *S*(*t, f*) over all frequencies:

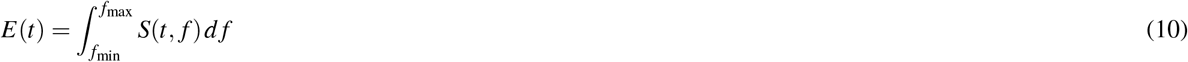

This integration process aggregates the spectral energy across different frequency bands, providing a comprehensive measure of the overall energy content of the audio signal at each time step *t*. By finding the peaks of these energy functions, we could identify significant increases in energy corresponding to the onsets of musical notes within the melody. Let *𝓅* be the set of time indices where *E*(*t*) has local maxima. These peaks served as reliable indicators of note onsets, allowing for precise localization of the timing at which each note begins:

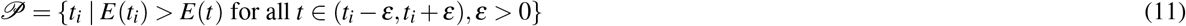

where *ε* is a small positive value used to define the neighborhood around each peak. Thus, the times *t*_*i*_ ∈ *𝓅* indicate the note onsets within the melody (Fig. 5).

### Babbling and Training the Inverse mapping

Since the system has no prior information on its dynamics, the physics of the environment, topology, or structure, it begins by exploring in a general sense through the execution of random control sequences to the motors, a process we refer to as motor babbling. After 2 mins. of motor babbling, the system uses the babbling data and creates the inverse map.

### Motor babbling

During this phase, the system executes random control sequences and collects the resulting melody played by the musician hand. The MLP and CNN used in the inverse mapping is then trained with this input-output pair to create an inverse map between the system inputs (motor activation levels) and desired system outputs (melody’s spectrogram).

### Random activation values for motor babbling

The motor activations (control sequences) during motor babbling were generated employing two pseudo-random number generators with a uniform distribution. The first generator randomly determines the activation order for each of the four motors (and, consequently, the keys). In contrast, the second generator selects the activation level between 75% and 100%.

### Structure of the ANN

Two neural network architectures were developed to extract the musical percepts from the spectrogram of the melodies: a CNN and an MLP. The CNN architecture consists of two convolutional layers with 32 and 64 filters, each followed by a max-pooling layer for downsampling. Subsequently, the flattened data is passed through two dense layers, each with 64 neurons and ReLU activation functions, before reaching the output layer with four neurons and a softmax activation function. In contrast, the MLP architecture comprises a single dense layer with 32 neurons and ReLU activation function, followed by the output layer with four neurons and softmax activation. Both models were compiled using the Adam optimizer and categorical cross-entropy loss function. The CNN model was trained for five epochs, while the MLP model was trained for twenty epochs, with validation data used to monitor accuracy. The architectures were implemented using the TensorFlow and Keras libraries in Python. Additionally, before training, the spectrogram data was normalized by dividing by the maximum absolute value using NumPy.

### Ethical Approval

All procedures were approved by the University of Southern California internal review board (USC IRB: HS-17-00304), and written consent was obtained from all participants prior to participation. The study conformed to all standards the Declaration of Helsinki set, except for registration in a database.

### Study Participants

Nine adults participated in this study. Five participants had no prior training in piano playing. Four participants had the ability to play the piano fluently.

### Experimental Setup

In this study, we used a four-finger tendon-driven hand, and a DC motor akin to a muscle provided the mechanical power to the joints. Each finger is driven by an individual tendon actuated by a DC brushless motor. The tendons are designed to passively return to their initial positions with the aid of springs. At the same time, the connection between the muscles and joints, analogous to the tendons in biological systems, is represented by a string referred to as a tendon within our robotic hand. Our modular tendon-driven robotic hand, equipped with DC motors, was managed by a custom DAQ and control program on a DAQ-capable computer with the NI-DAQmx driver. Operating on a machine learning-capable computer, our perceptual learning pipeline utilized RT-Bridge over Ethernet to transmit commands and receive sensory data from the DAQ program with a sub-millisecond round-trip latency. to play and record the melodies, we use a MIDI device connected to a computer that runs GarageBand, and then the recorded files were exported as wave files to be used by the algorithm in Python.

### Melodies

For our study, three original melodies were meticulously composed for this research (Figure 7). These melodies were deliberately crafted to utilize only four distinct keys: F3, E3, D3 and C3. Each melody was performed at a consistent tempo of 90 beats per minute, executed by our two skilled composers on a Steinway grand piano. This selection of keys and tempo ensured a controlled and uniform musical environment, facilitating precise analysis and comparison within our experimental framework.

## Author Contributions

### Funding

Funding: NSF CRCNS Japan-US 2113096; DARPA L2M Program W911NF1820264; Division of Biokinesiology and Physical Therapy Graduate Teaching Assistantships.

## Acknowledgments

We had the privilege of collaborating with two composers, Richard Tuttobene and Targol Karimi Moghaddam, who graciously crafted three new melodies. These pieces capture melodious musical dynamics using only four adjacent notes.

